# Common neural responses to narrative speech in disorders of consciousness

**DOI:** 10.1101/166405

**Authors:** Ivan Iotzov, Brian C Fidali, Agustin Petroni, Mary M Conte, Nicholas D Schiff, Lucas C Parra

## Abstract

**Objective:** Clinical assessment of auditory attention in patients with disorders of consciousness is often limited by motor impairment. Here, we employ inter-subject correlations among electroencephalography responses to naturalistic speech in order to assay auditory attention among patients and healthy controls.

**Methods:** Electroencephalographic data were recorded from 20 subjects with disorders of consciousness and 14 healthy controls during of two narrative audio stimuli, presented both forwards and time-reversed. Inter-subject correlation of evoked electroencephalography signals were calculated, comparing responses of both groups to those of the healthy control subjects. This analysis was performed blinded and subsequently compared to the diagnostic status of each patient based on the Coma Recovery Scale-Revised.

**Results:** Subjects with disorders of consciousness exhibit significantly lower inter-subject correlation than healthy controls during narrative speech. Additionally, while healthy subjects had higher inter-subject correlation values in forward vs. backwards presentation, neural responses did not vary significantly with the direction of playback in subjects with disorders of consciousness. Increased inter-subject correlation values in the backward speech condition were noted with improving disorder of consciousness diagnosis, both in cross-sectional analysis and in a subset of patients with longitudinal data.

**Interpretation:** Inter-subject correlation of neural responses to narrative speech audition differentiates healthy controls from patients and appears to index clinical diagnoses in disorders of consciousness.

## Introduction

Patients with chronic disorders of consciousness (DOC) have varied outcomes that are difficult to prognosticate^1, 2^. Accurate assessment of higher-level cognitive abilities such as auditory attention is essential for accurate diagnosis and may determine candidacy for assistive communication devices. However, many patients have impaired channels of motor communication, resulting in a mismatch between the clinical assessment of auditory comprehension and neuroimaging evidence^3, 4^. Thus, quantifying auditory attention in this population is an urgent research priority.

Metabolic studies have demonstrated a difference in cortical auditory processing between vegetative state (VS) patients that lack the ability to interact with their environment and minimally-conscious (MCS) patients, who demonstrate behavioral interactions^5, 6, 7, 8^. Attempts to discriminate auditory attention and processing in these MCS patients with electroencephalography (EEG) have focused on event-related potential paradigms utilizing single words and repeated sound sequences^9, 10, 11, 12, 13^. While such EEG measures appear to index cognitive processes, they are unable to assess the presence of sustained auditory attention that is required for patients to comprehend speech in everyday environments.

Here, we investigate the presence of sustained auditory attention in MCS patients by measuring neural responses to narrative speech. In healthy populations, prolonged attention to a stimulus results in entrainment of the subject’s evoked activity not only to low-level features of the stimulus itself, but also the cortical activity of other subjects experiencing the same stimulus^14, 15, 16, 17^. This effect has been observed in visual as well as auditory contexts for EEG^18, 15, 16^, functional magnetic resonance imaging^19, 20^, and magnetoencephalography^21, 22^. Moreover, inter-subject correlation (ISC) of EEG evoked responses has been shown to discriminate attention better than conventional EEG measures^15^ and is able to predict selective auditory attention during auditory stream segregation^23, 17^.

In this single-blinded study, we employ inter-subject correlation of EEG to assess sustained auditory processing of narrative speech. Twenty patients with disorders of consciousness and 14 healthy controls were presented two narratives in both, a forwards and a backwards (time-reversed) condition. We predicted that patients will have lower ISC of the EEG evoked activity as compared to healthy control subjects. ISC for healthy group and patients were extracted from time-locked EEG without knowledge of individual patient diagnoses. Individual subjects’ ISC scores for forward and backwards speech presentation were compared healthy controls and among clinical diagnoses. Finally, we discuss the relevance of our findings in the search for biomarkers of auditory attention among disorders of consciousness.

## Methods

### Subject recruitment

Healthy control and DOC subjects included were drawn from a convenience sample available from a multi-day, inpatient hospital admission research study approved by the Weill Cornell Medical College Institutional Review Board. The study collects video electroencephalography (EEG), multimodal neuroimaging data, and clinical outcome measures in the context of chronic disorders of consciousness resulting from severe acquired brain injury. Healthy control subjects provided written consent, while consent was obtained from the legally authorized representatives of the DOC subjects.

### Clinical outcome measures & blinding

Clinical assessments were made by serial administrations of the Coma Recovery Scale-Revised (CRS-R)^24^ by neurologists during inpatient research admissions. Sub-scale scores from each patient’s highest CRS-R and command following data from functional neuroimaging were assessed by an expert neurologist (senior author NS) to codify clinical diagnoses of VS, MCS-/+, or eMCS according to the the following criteria: Patients may remain wakeful but unresponsive to the external world in the persistent vegetative state (VS)^25^. Others in the minimally conscious state (MCS) may have inconsistent responses to their surroundings^26^. This category is subdivided into plus/minus, with MCS- designating individuals with exclusively low-level, reflexive behavior such as withdrawing from pain or turning towards sound. In contrast, the presence of higher-level cognitive functions like inconsistent command following, yes/no questions, or intelligible vocalizations earns a designation of MCS-plus (MCS+)^27^. Patients emerged from the minimally conscious state (eMCS) can interact with their surroundings through functional object use or communicate reliably. Authors II, AP & LP were blinded to these clinical diagnoses, with an agreement to un-blind clinical diagnoses after inter-subject correlation scores were generated for all individuals across all visits. Health status (healthy control versus DOC patient) was not blinded in order to facilitate development of the ISC metric, which compares patients to healthy controls.

### Stimulus presentation

A female narration of Lewis Carroll’s *Alice in Wonderland* audio file (148 sec) was converted to 44100 kHz WAV file for the *Alice* stimulus. It was subsequently reversed in time using Audacity (audacity.sourceforge.net) to create the backward *Alice* stimulus. Forward and backwards audio for a live performance of a stand-up comedy with music interlude, *Pieman* (Jim O’Grady; length 7 min), were obtained from Uri Hasson^28^.

With either stimulus, subjects were instructed to listen carefully to the story that they were to hear through their headphones. Both forward and backward audio files were then played with Presentation software (Neurobehavioral Systems, Inc), with mono audio presented binaurally through Etymotic Research ER3A earphones. Audio start and stop points were time-locked into the video EEG record using photic stimulation markers in the Natus Neuroworks software and paradigm audio was also recorded along with the video. One iteration each of forward and backward audio were interleaved with 30 seconds of rest (*Alice;* Forwards-Rest-Backwards) or 60 seconds of rest (*Pieman;* Backwards-Rest-Forwards).

DOC subjects participated in a 2-3 day overnight study while HC subjects participated in a 24 hour study. During each subject’s study, the *Alice* paradigm was repeated 2-6 times in an effort to ensure at least one block of stimulus presentation with limited artifact. Of the healthy controls, 10 repeated the same paradigms at a six-month revisit. Additionally, four patients had a second visit 1-3 years after their initial visit, three with *Alice* data. *Pieman* data were presented once per subject visit and not repeated upon subject re-visit. All stimuli were presented while subjects were in an eyes-open, wakeful state.

### Data collection

The EEG data were recorded using 37 electrodes (Nihon Kohden (Japan) silver-collodion disc electrodes, 10 mm) placed via an enhanced 10-20 arrangement, using the Natus XLTEK (Oakville, Canada) system. EEG was recorded with synchronized video. The typical inter-electrode spacing was 3 to 4 cm and impedances were maintained ≤ 5 kΩ. Bipolar referencing was used, with a FCz reference and AFz ground electrode. Bilateral electrooculography (2 leads) and electrocardiography were also recorded. Signals were amplified and digitized at 250 Hz using an anti-aliasing high-pass filter with a corner frequency at 0.4 times the digitization rate.

### Data extraction and export

Video EEG data were reviewed in Natus Neuroworks software. *Alice* & *Pieman* trials in which subjects remained in an eyes-closed state for over 10 consecutive seconds or exhibited sustained vocalization and movement were not considered for further analysis. The remaining paradigms were exported to ASCII text files and imported to MATLAB (8.3). Forward and backwards conditions were exported using in-house scripts for subsequent analysis.

In the *Alice* stimulus set, data from 14 healthy controls—10 with two visits at a 6 month latency—were submitted for analysis for a total of 48 datasets. Data from 16 patient subjects (3 longitudinal) totalled 55 datasets. Of these subjects, In the *Pieman* stimulus set, data from 12 healthy controls from single visits (one longitudinal) were submitted, yielding a total of 13 healthy control datasets. Data from 20 patient subjects (four with longitudinal data) were submitted for analysis, for a total of 22 datasets. Identifiers for patients vs controls were coded before submission to author II for single-blinded analysis.

Upon visual inspection, and prior to processing, some data had to be discarded due to limited quality, excessive movement artefacts, or a large number of missing electrodes (5 more more channels). For the *Alice* stimulus, we are left with 13 healthy controls (2 of whom were only presented the *Alice* stimulus) for a total of 43 repeats and 11 DOC patients (1 of whom was only presented the *Alice* stimulus) for a total of 38 repeats, and a grand total of 81 repeats for Alice. For the *Pieman* stimulus, we analyzed 12 healthy controls (1 unique to *Pieman*) with a total of 13 repeats, as well as 19 DOC patients (9 unique to *Pieman*) with 22 repeats, and a grand total of 35 repeats.

### Preprocessing

Data analysis follows Ki et al., 2016. Briefly, for each stimulus, raw EEG data was epoched across repetitions and filtered to remove drift and power-line noise (0.5Hz 5th order high-pass and a 60Hz 10th order band-stop Butterworth filter respectively; extra padding of 2 seconds in each epoch was removed after filtering). Eye movement artefacts were removed by regressing out activity from two EOG electrodes and Fp1 from all EEG electrodes. Finally, we used robust principal component analysis (rPCA)^29^ as an artifact rejection procedure with regularization parameters λ = 0.0052 for *Alice* and λ = 0.003 for *Pieman* data. This procedure effectively interpolates outlier activity based on the principal subspace of the data.

### Inter-subject correlation analysis

Our previous work suggests that attention to ongoing narrative speech can be reliably measured by correlating the evoked EEG of an individual subject to that of an attentive group, with more attentive subject exhibiting higher inter-subject correlation (ISC)^15^. ISC is best measured, not on individual electrodes, but on the components of EEG that maximally correlate across subjects^30^ (we used code published in Cohen 2016 for these computations). Here correlated components were optimized on data available for the *Alice* stimulus including DOC patients and healthy controls in both the forward and backward conditions. This gives us components with spatial distribution (Fig. 1A) resembling results from previous work on auditory stimuli^18, 15^. Spatial distributions for the *Pieman* data did not replicate previous results indicating that these data were not sufficient for component extraction (smaller and noisier sample compared to previous work). However, using the components extracted from *Alice*, we could compute ISC for both *Alice* (Fig. 1B) and *Pieman* stimuli (Fig. 1C). Here we measure ISC for each subject by correlating component activity to that of the healthy control group, averaging over all possible pairings that involve a given subject. As in previous work, we use the sum of the ISC values computed in the first 3 strongest components. ISC was computed for each recording, and then averaged across all available repetitions for a given subject. For Figure 1 this was further averaged across subsequent visits, where available.

**Figure 1.**
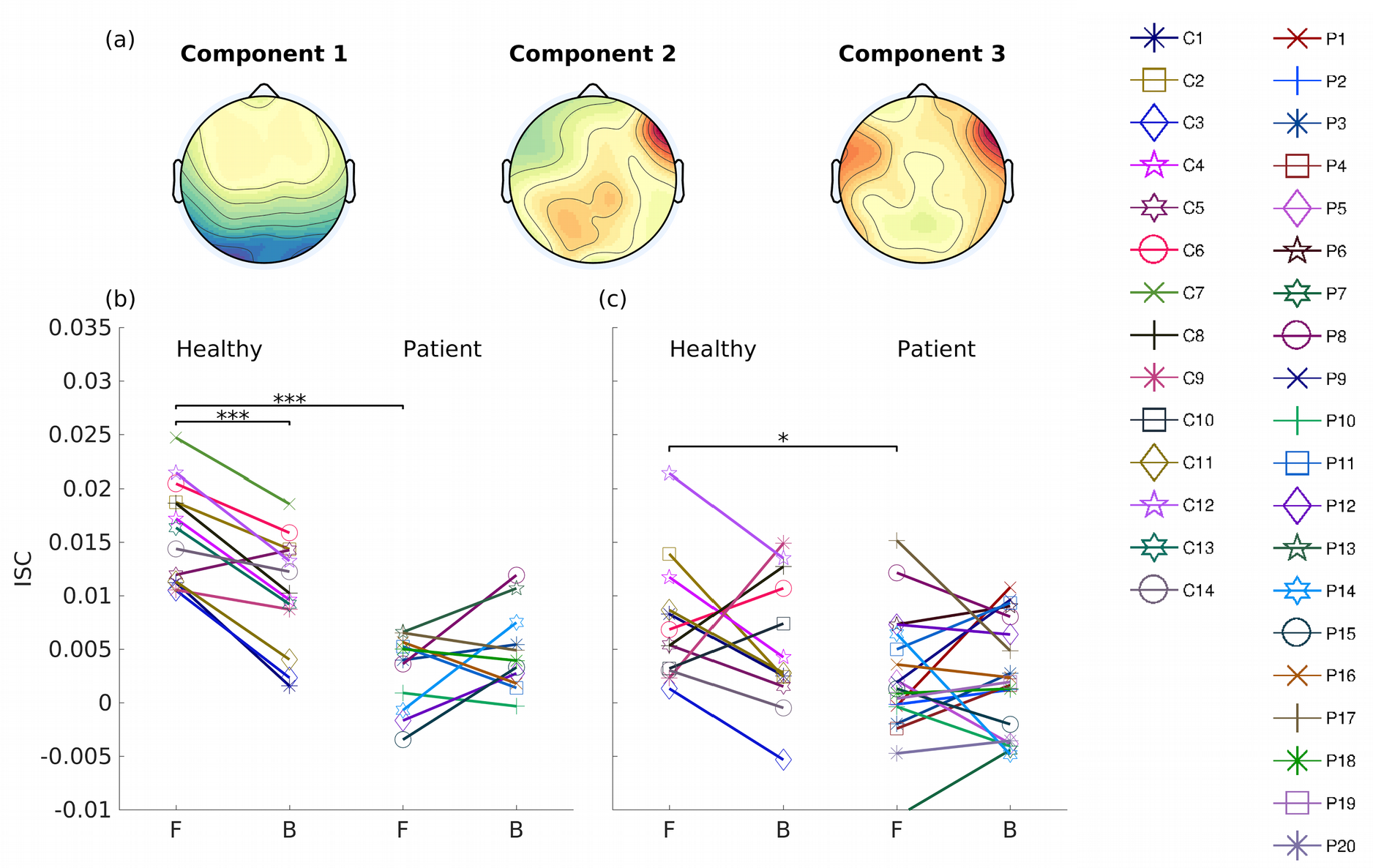
Inter-subject correlation of EEG responses evoked by auditory narratives in DOC patients and healthy controls. **(a)**Spatial components of correlated activity between subjects. These three components capture the strongest inter-subject correlation (ISC) and were computed here over all conditions in both patients and healthy controls using “Alice” -- a segment of “Alice in Wonderland” narrated by a female speaker (148s long). **(b)** ISC of healthy controls (N=13) and patients (N=11) during the *Alice* stimulus. ISC is measured by correlating component activity of each subject to the cohort of healthy controls and summing over the first 3 components. It is measured separately for Forward (F) and Backward (B) conditions and averaged over repeated renditions and visits. **(c)** Same as in panel (b) but for “Pieman” -- a 6 min live recording of a stand-up comedy performance for healthy controls (N=12) and patients (N=19). Significant post-hoc pairwise comparison are shown as black horizontal lines (*** p<0.001, ** p<0.01, * p<0.05, uncorrected).

## Results

### EEG inter-subject correlation during auditory narratives is reduced in DOC patients as compared to healthy controls

Fourteen healthy controls (39 +/- 11 years of age, 8 men) and 20 disorders of consciousness (DOC) patients (31 +/-13 years of age, 4 women) were included in the analysis. Video EEG were recorded while subjects were presented audio narratives through headphones. In the first experiment, subjects heard a short segment from a professionally narrated audiobook of *Alice in Wonderland* (148 sec). This was presented to 13 healthy controls (2 unique to the *Alice* stimulus), and 11 DOC patients (1 unique to the *Alice* stimulus). The sound was also played back time-reversed, resulting in a forward and backward playback, which was repeated several times for each subject (N = 3.3 ±1.4 for controls and N=3.5 ±1.8 for patients). Some subjects participated in two visits on separate days, resulting in a total of 81 recordings included in subsequent analysis (see Methods 2.3).

We extracted components of the EEG that were maximally correlated between subjects during presentation of the *Alice* stimulus following established procedures^18, 30^ (see Methods 2.7). Figure 1A shows the three correlated components that capture most of the inter-subject correlation (ISC) in these data. These are consistent with previous findings for auditory narratives^18, 15^. We measured ISC in these three components, correlating both patients and healthy participants to the healthy participants. Thus, ISC measured for each subject, how similar evoked responses are to those of a healthy normative group. ISC values are shown in Figure 1 for each subject. As expected, patients have lower ISC compared to the control group in particular for the *Alice* stimulus (Fig. 1B). Furthermore, backward playback reduced ISC, at least in health participants.

To test for statistical significance of these effects we performed a two-way ANOVA with fixed factors of playback conditions (forward/backward) and health status (control/patient). To control for the evident variability of ISC across subjects we included subject identity as a random effect. We found a strong effect for health status (F(1, 114) = 32.76, p = 5.4x10^-6^) and a strong interaction between playback condition and health status (F(1, 114) = 18.23, p = 1.3x10^-4^). The random effect of subject was highly significant (F(22, 114) = 4.05, p = 8.8x10^-4^) indicating that ISC is quite variable across subjects. Follow-up comparison shows that ISC drops for backward playback in healthy controls (t(12) = 5.97, p = 6.5x10^-5^). ISC is also lower for patients as compared to controls in the forward playback condition (t(22) = 7.59, p = 1.4x10^-7^).

A more limited dataset was also available for an audio narrative involving a live recording of standup comedy (*Pieman*, Jim O’Grady)^28^, in forward and backward playback (12 healthy controls and 19 DOC patients). Unlike the *Alice* stimulus, only one recording was available per subject in healthy controls and patient datasets (resulting is a total of 35 recordings). ISC was computed for this data set using the same components extracted with the *Alice* stimulus (Figure 1C). As with the *Alice* datasets, a two-way ANOVA was performed to test for differences in ISC with fixed factors of health status and playback condition, and subjects as random factor. We find again a contrast between patients and healthy participants (F(1, 8) = 5.52, p = 0.0255), driven by a contrast in the forward condition, (t(29) = 2.489, p = 0.019) but this time no effect from the playback condition (F(1, 8) = 0.62, p = 0.44) and no interaction between playback condition and health status (F(1, 8) = 0.9, p = 0.35). A three way ANOVA with stimulus as additional factor (*Alice* vs *Pieman*) confirms that the *Pieman* story elicited overall lower ISC values (F(1, 142) = 9.24, p = 0.006).

### ISC during backward playback of speech correlates with diagnostic status of DOC patients

The previous analysis was done blinded to the clinical diagnosis of the patients. Patients had suffered a variety of etiologies and carried one of four diagnoses: vegetative state (VS), minimally conscious state minus (MCS-), MCS plus (MCS+), and emerged from MCS (eMCS; see Section 2.2). After unblinding, these diagnoses were coded as a categorical variable for statistical analysis. We performed three planned comparisons between this categorical diagnosis and ISC, namely, forward and backward conditions as well as their difference as a possible control for the evident variability in ISC across subjects.

Figure 2 shows the comparison of forward and backwards presentation ISC values using the *Alice* data across these diagnosis groups MCS–, MCS+, and eMCS. Here we separated visits 1 and 2 as the diagnostic criteria also changed for the patients where data from two visits were available. The single patient in vegetative state (P7) did not have an *Alice* recording, so this diagnosis group was not presented. A one-way ANOVA with diagnostic state as factor shows a significant effect for ISC backward presentation (F(2, 11)=9.46, p=0.0041) which remains significant after correction for the 3 planned comparisons (p=0.012). Figure 2(b) suggest that this effect results from an increase of ISC for backward speech with improving diagnosis across patients.

**Figure 2.**
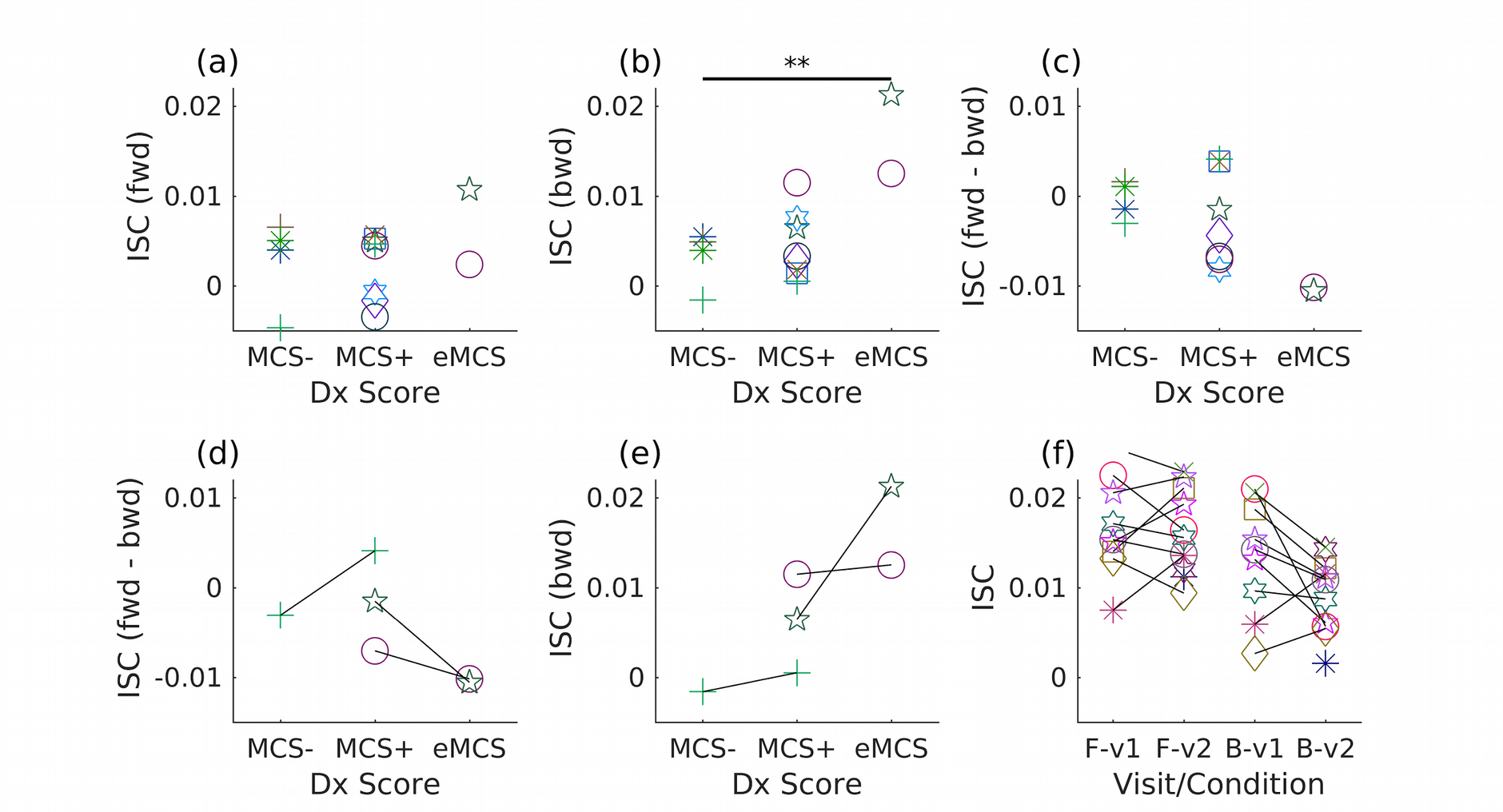
Comparison of ISC with clinical diagnosis in DOC patients. (a) ISC for the *Alice* stimulus during forward playback and (b) backward playback for N=11 patients. Black line represents significant influence of diagnosis on ISC score (p < 0.01) (c) difference in ISC between forward and backward playback. Here visit 1 and 2 are separated as diagnosis also changes across visits. (d) Change in ISC difference over two visits for the three patients for which this data was available. (e) as in panel (d) but for backward playback. (f) for reference, we show here the variability of ISC measures across visits in healthy controls. No data for the *Alice* condition, was available from vegetative state (VS) patients in this sample. Subject symbols and colors are consistent with Figure 1.

For three patients two time points of recordings for the *Alice* stimulus were available separated by 12 months (P8), 36 months (P10), and 17 months (P13). During this time, diagnostic score improved on all 3 subjects (Table 1). For all three patients ISC to the backward speech increased along with the clinical diagnosis (Fig. 2E), although the change appears meaningful for only one of them, considering the normal fluctuations seen in healthy controls across visits (Fig. 2F). For this patient, brain metabolism as measured with positron emission tomography (PET) markedly increase from visit 1 to visit 2 (Supp. Fig. 1). The clinical history of this patient is described in detail in the supplemental text.

**Table 1.** Patient Demographics and diagnoses: Demographics of all 20 disorders of consciousness (DOC) subjects included in this study. Age at time of study as well as age of acquired brain injury are reported in years. Documentation of the Coma Recovery Scale-Revised (CRS-R) and its subscales are as previously reported (Section 2.2). Abbreviations: VS = vegetative state; MCS = minimally conscious state; eMCS = emerged from minimally conscious state; W = white/Caucasian, NH = non-Hispanic, TBI = traumatic brain injury; HAI = hypoxic/anoxic injury; CA = cardiac arrest; SAH = subarachnoid hemorrhage. * denotes emergence from MCS, $ full case report in Forgacs et al., 2016^32^.

| Patient code | Visit age | Injury age | Sex | Race | Injury type | Total CRS-R | Auditory (0-4) | Visual (0-5) | Motor (0-6) | Oro-motor (0-3) | Communication (0-2) | Arousal (0-3) | CMD | Diagnosis |
| --- | --- | --- | --- | --- | --- | --- | --- | --- | --- | --- | --- | --- | --- | --- |
| P1 | 45 | 35 | F | W (NH) | encephalitis | 19 | 4 | 4 | 6* | 2 | 1 | 2 |  | eMCS |
| P2 | 30 | 23 | M | W (NH) | TBI | 23 | 4 | 5 | 6* | 3 | 2* | 3 |  | eMCS |
| P3-v1 | 24 | 16 | M | W (NH) | TBI | 9 | 2 | 2 | 3 | 0 | 0 | 2 |  | MCS- |
| P4 | 57 | 53 | M | W (NH) | TBI | 21 | 4 | 5 | 5 | 3 | 1 | 3 |  | MCS+ |
| P5 | 20 | 17 | F | W (NH) | HAI (CA) | 6 | 1 | 0 | 1 | 2 | 0 | 2 | Yes (EEG) \$ | MCS+ |
| P6 | 40 | 37 | M | W (NH) | HAI | 15 | 4 | 3 | 2 | 3 | 1 | 2 |  | MCS+ |
| P7 | 23 | 20 | M | Asian | TBI | 6 | 2 | 0 | 1 | 1 | 0 | 2 |  | VS |
| P8-v1 | 27 | 22 | M | W (NH) | TBI | 17 | 4 | 3 | 6* | 1 | 1 | 2 |  | eMCS |
| P8-v2 | 28 |  |  |  |  | 14 | 4 | 2 | 5 | 1 | 0 | 2 |  | MCS+ |
| P9 | 36 | 19 | M | W (NH) | TBI | 19 | 4 | 4 | 6* | 2 | 1 | 2 |  | eMCS |
| P10-v1 | 23 | 12 | F | W (NH) | TBI | 12 | 2 | 3 | 3 | 2 | 0 | 2 |  | MCS- |
| P10-v2 | 26 |  |  |  |  | 13 | 4 | 3 | 2 | 2 | 0 | 2 |  | MCS+ |
| P11 | 21 | 17 | M | W (NH) | HAI | 16 | 4 | 2 | 4 | 3 | 1 | 2 |  | MCS+ |
| P12 | 25 | 20 | M | W (NH) | TBI | 17 | 4 | 4 | 4 | 3 | 0 | 2 |  | MCS+ |
| P13-v1 | 19 | 18 | F | Black | TBI | 10 | 3 | 3 | 2 | 0 | 0 | 2 | Yes (fMRI) | MCS+ |
| P13-v2 | 20 |  |  |  |  | 23 | 4 | 5 | 6* | 3 | 2* | 3 |  | eMCS |
| P14 | 26 | 23 | M | Black | TBI | 11 | 2 | 3 | 3 | 1 | 0 | 2 | Yes (fMRI) | MCS+ |
| P15 | 57 | 54 | M | W (NH) | SAH | 19 | 4 | 5 | 5 | 2 | 1 | 2 |  | MCS+ |
| P16 | 22 | 21 | M | W (NH) | TBI | 17 | 4 | 4 | 4 | 2 | 1 | 2 |  | MCS+ |
| P17 | 26 | 25 | M | W (NH) | TBI | 6 | 1 | 3 | 1 | 0 | 0 | 1 |  | MCS- |
| P18 | 21 | 19 | M | W (NH) | TBI | 10 | 1 | 3 | 3 | 1 | 0 | 2 |  | MCS- |
| P19 | 56 | 55 | M | W (NH) | HAI (CA) | 21 | 4 | 5 | 5 | 3 | 1 | 3 |  | MCS+ |
| P20 | 23 | 19 | M | W (NH) | TBI | 5 | 4 | 0 | 0 | 0 | 1 | 0 | Yes (fMRI) | MCS+ |

## Discussion

To our knowledge, this is the first study to assay electroencephalographic responses to naturalistic speech in patients with disorders of consciousness (DOC). By correlating EEG responses to those of healthy controls during auditory narratives, we demonstrate that healthy controls have more similar within-group responses than do patients. The contrast between forward and backward speech observed in healthy controls is absent in patients, although the strength of response to backward speech appears to be linked to diagnostic score.

The topography of the three most salient inter-subject correlation (ISC) components and higher forward vs. backwards ISC scores in our healthy controls (Fig. 1) closely replicate prior studies with narrative speech^18, 15^. In healthy populations, we interpret ISC as a measure of the reliability of auditory evoked responses, which are modulated by attention. As ISC scores in narrative speech is modulated by attention^15^, the DOC patients’ lower ISC scores could be interpreted as evidence of impairment of normal auditory attention. Additionally, the heterogenous injury patterns of DOC patients might independently contribute to this effect, as ISC was computed here by correlating EEG of patients with that of healthy controls.

In addition to having lower ISC values than healthy controls, the contrast between forwards and backwards ISC scores was absent in this group of DOC patients (Fig. 1B-C). Interestingly, the absolute value of the backward ISC scores correlated positively with with clinical diagnosis (Fig. 2B). This finding contrasts with a prior functional magnetic resonance imaging (fMRI) study with naturalistic speech, where time-reversed stories elicited stronger blood-oxygen level dependent (BOLD) responses in controls than forwards presentation, and two minimally conscious state patients lacked BOLD responses entirely to backwards stimuli^8^. These divergent group findings in controls could reflect methodology; EEG recordings have much higher temporal resolution than BOLD signals, directly reflect neuronal activity, and robustly encode lower-level auditory features. Perhaps, ISC might reflect the novelty and perceptual salience of backwards stimuli in patients, but are less attended to by healthy individuals who quickly recognize the stimulus as non-speech, and therefore lose interest. Regardless, as the strength of backwards ISC response correlates positively with clinical diagnosis and reliable auditory attention is required for MCS+ and eMCS diagnoses, ISC values appear to index a clinically important element of auditory processing.

Our study has several limitations. Primarily, the sample size of 20 DOC patients, with only 12 that could be compared to the clinical diagnosis and only a single case of vegetative state in our convenience sample. This precludes validation of this measure as a diagnostic tool, despite significant differences in backwards ISC across diagnoses. The more variable ISC values in the *Pieman* dataset, despite longer stimulus length, are primarily attributed to the single recording available for each subject. The higher ISC scores for *Alice*, may be the result of the more clearly-articulated studio recording as compared to the live recording of the *Pieman* stimulus. Lastly, these paradigms were presented as part of a rigorous testing schedule throughout overnight visits; such a schedule would be expected to reduce ISC values, as ISC scores decrease after repeated trials in healthy individuals^15^. Future studies would benefit from more stimulus repetitions during outpatient testing to minimize fatigue and improve the reference dataset among healthy control participants.

While our data establish ISC of neural activity during audition as a possible biomarker for auditory processing in DOC, interpretative caution is required. For example, Patient 13 demonstrated remarkable clinical improvement, which correlated with an increase in ISC values in the second visit (Figs. 2D, 3 [green star]; Supplement). While these findings are encouraging and mirror her recovery of communication, we argue against inferring the quality of narrative capacity or subjective conscious experience of this or any individual patient^31^, as neither EEG nor BOLD signals can be causally linked to higher cognitive functions. Instead, we focus on the diagnostic and prognostic promise of neurophysiological measures like EEG responses to narrative speech.

In summary, we present the first evidence that EEG evoked responses to narrative speech in DOC patients may reflect clinically important elements of auditory processing. Further research is needed to untangle the cognitive processes required for higher-level attention and cognition from the cortical markers of more basic auditory processing. That said, these data demonstrate the potential of correlating neural activity in response to naturalistic speech to that of healthy controls as an index of auditory processing that might be developed into a diagnostic tool in the search for covert cognition in these patients.

## Acknowledgement

We are indebted to Jaco Sitt for his helpful discussions, Ofer Tchernichovski for proposing this study, and Uri Hasson for sharing his stimulus files. We thank Esteban Fridman for contributing Supplemental Figure 1. We also thank Tanya Nauvel and Henning Voss for insightful comments as well as Zoe Adams & Billy Curley for data collection. BF was supported by NIH/NCATS #TL1TR000459, and NS by NIH #5R01HD51912, the Jerold B. Katz Foundation, & the James S. McDonnell Foundation.

## Author contributions

BF and LP designed the study. BF and MC collected patient data with subjects recruited by NS. BF reviewed and exporting video EEG data. AP developed analysis codes and conducted the pilot data analysis. II conducted advanced preprocessing as well as primary and secondary analyses, under the direction of LP. BF, II, & LP wrote the manuscript with contributions from MC & NS.

## Supplemental Material

### Patient 13 clinical history

Patient 13 was a previously a healthy woman before suffering a severe traumatic brain injury as an unrestrained passenger in a motor vehicle accident at age 18. After emergent bolt craniotomy and exploratory laparotomy, she had a prolonged medical course before extubation and transfer to rehabilitation facility. Upon emergence from coma, Patient 13 was wakeful and demonstrated minimal interactiveness. Family members reported periods of increased lucidity and command following during the following months. Amantadine was added without clear benefit.

On her first research visit (17 months after injury), Patient 13 would localize to sound and withdraw from painful stimuli in all extremities. She demonstrated consistent sleep/wake cycles, but was often somnolent in the absence of external stimulation during the day. Relevant medications included baclofen 20 mg three times daily for spasticity, amantadine 200 mg twice daily, and levetiracetam 500 mg twice daily for seizure prophylaxis (no known seizure history). Her total Coma Recovery Scale-Revised (CRS-R) scores of 9-10 (see Table 1), meeting criteria for MCS– state. In light of positive fMRI command following, she was considered in a state of cognitive-motor dissociation, meriting a final diagnosis of MCS+.

On her second visit (34 months after injury), Patient 13’s neurological exam improved dramatically. She was now regarding and visually tracking consistently, forming intelligible verbalizations including yes/no answers, and manipulating objects such as a hairbrush appropriately. Her total CRS-R scores were now 21 & 22 on repeated testing (see Table 1), and Mississippi Aphasia Screening Test (MAST; Nakase-Thompson, 2004) scores were 46 & 47 out of 50 on repeated testing. Her second visit diagnosis was emerged from MCS (eMCS), with her behavioral assessment most consistent with a confusional state.

Between visits, metabolic imaging (18F-FDG-PET; Supp. Fig 1) also demonstrated diffusely increased metabolism, most prominent in bilateral temporal and occipital lobes as well as L>R thalamus. In both visits, her electroencephalographic record demonstrated anterior-posterior gradient with a posterior dominant rhythm approaching 8-9 Hz, but not consistently maintained. Periods of right>left focal slowing were noted, and epileptogenic features absent.

#### Sources

Nakase-Thompson, R. (2004). The Mississippi Aphasia Screening Test. The Center for Outcome Measurement in Brain Injury. http://www.tbims.org/combi/mast (accessed June 27, 2017).

---

**Supplemental Figure 1:**
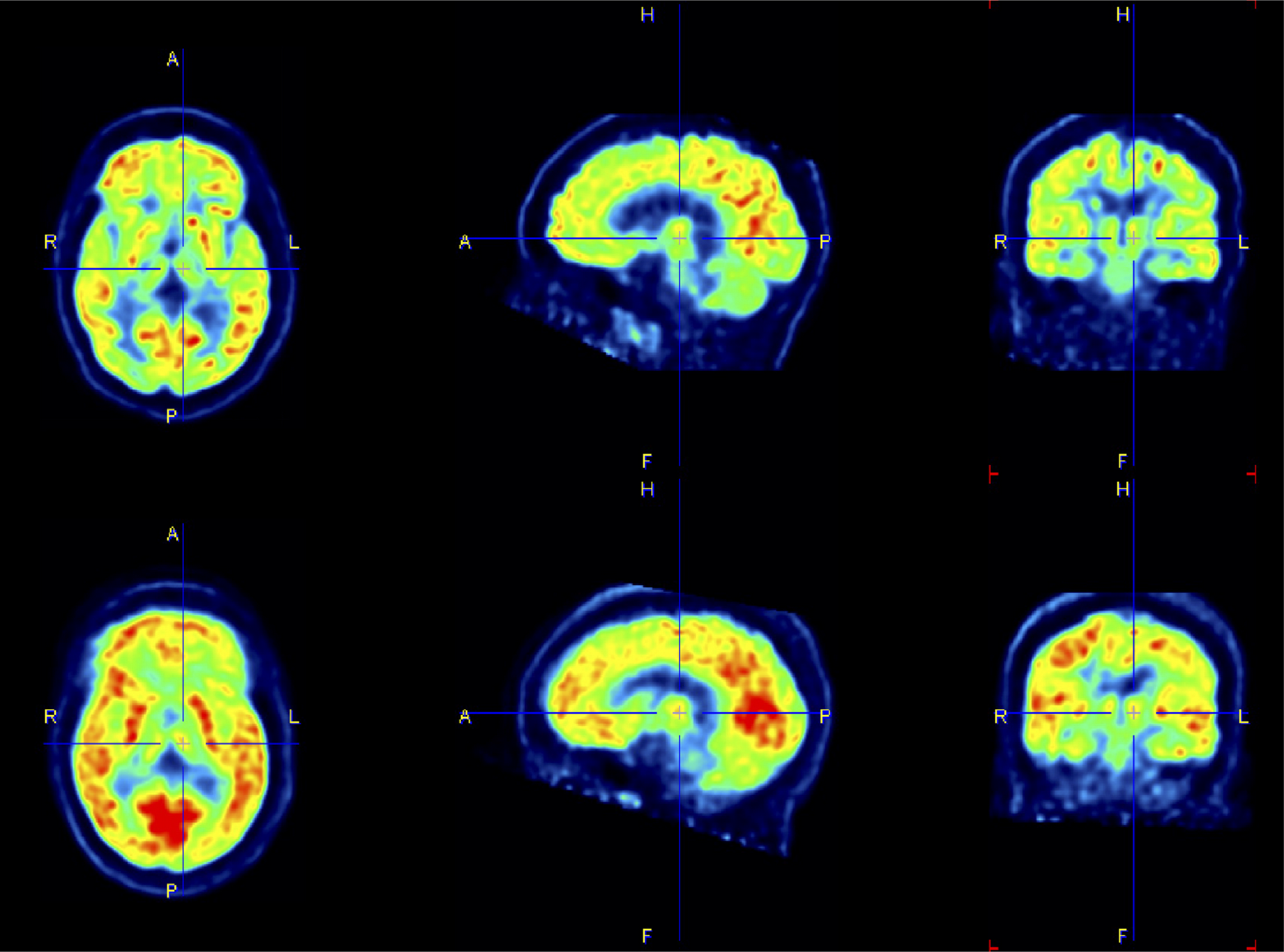
Interval increases in metabolic activity mirror clinical improvement in subject P13. Global metabolism (measured here in 18F-FDG-PET uptake) increases from Visit 1 (top row) to Visit 2 (bottom row) in axial (L column), R para-sagittal (middle column), and coronal (R column) sections. Coordinate on L central thalamus. Standardized uptake values (SUV)were normalized between visits by weight. Changes are most prominent in bilateral temporal, parietal, and occipital cortices. Increased uptake is seen as well in the basal ganglia, prefrontal cortex, and L>R thalamus.

